# Expanding the Frontiers of Structural Analysis in Short RNAs by Ultra-High Field 1.3 GHz NMR

**DOI:** 10.64898/2026.08.23.746555

**Authors:** Naoya Tochio, Taiichi Sakamoto, Takanori Kigawa

**Affiliations:** RIKEN Center for Integrative Medical Sciences, Kanagawa, Japan; Department of Life Science, Faculty of Advanced Engineering, Chiba Institute of Technology, Chiba, Japan

**Keywords:** Ultra-high field NMR, RNA aptamer, HIV-1 Viral infectivity factor, Residual Dipolar Coupling

## Abstract

Residual dipolar couplings (RDCs) obtained via magnetic field-induced alignment offer a powerful, media-free approach for the structural analysis of biomolecules. However, their detection in short, fast-tumbling nucleic acids remains elusive at conventional magnetic fields due to insufficient alignment and sensitivity. Here, we demonstrate the direct observation of these RDCs at 1.3 GHz in a 14-mer hairpin fragment derived from an HIV-1 Vif-targeting aptamer. The ^1^*J*_NH_ scalar couplings of imino protons were measured at fields ranging from 600 MHz to 1.3 GHz. While the coupling constants remained invariant between 600 and 900 MHz, a clear deviation was exclusively captured at 1.3 GHz for all base-paired stem residues, demonstrating the first media-free detection of field-induced RDCs in a short RNA of this size. This breakthrough arises from a synergistic 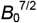 scaling, combining enhanced alignment 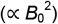 and sensitivity 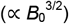. These RDCs showed excellent agreement with the NOE-derived structure. Additionally, the flexible loop residue G8 exhibited no detectable RDC, but displayed a field-dependent TROSY/anti-TROSY intensity inversion at 1.3 GHz, reflecting an unusual ^1^H chemical shift anisotropy (CSA) tensor that corroborates the local base-packing environment. Our findings highlight 1.3 GHz NMR as an indispensable tool for the structural analysis of short RNAs.

## Introduction

Residual dipolar couplings (RDCs) are a powerful tool in solution NMR spectroscopy. Because they provide long-range geometric information about the orientation of chemical bonds relative to the static magnetic field axis, RDCs are widely used for structural determination and validation of biomolecules.[1, 2] In an isotropic solution, dipolar interactions are averaged to zero by rapid molecular tumbling. Therefore, observing RDCs generally requires the induction of a weak alignment, typically achieved using external alignment media such as liquid crystals or alignment gels [3]. Alternatively, this weak alignment can be induced by exploiting the molecule’s intrinsic magnetic susceptibility anisotropy (Δ*χ*) in a strong magnetic field [4]. This field-induced alignment is proportional to the square of the magnetic field strength 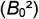, where the net alignment results from a balance between this magnetic energy and thermal fluctuation (*kT*).[1] [5] However, this effect is inherently subtle. For short nucleic acids and low-mass molecules, detecting field-induced alignment at conventional magnetic fields remains practically elusive.[2] [6] Recent advances in ultra-high-field NMR instruments above 1.2 GHz have significantly enhanced both sensitivity and resolution, providing the capability to resolve subtle structural and spectral variations.[7] This combined improvement in sensitivity 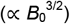 and resolution is vital for distinguishing precise changes in coupling constants.[8] Nevertheless, quantitative studies on the detection of field-induced RDCs in short RNAs, as well as their precise magnetic field dependence, remain limited.[2]

In this study, we performed precise measurements of ^1^H-^15^N couplings of imino protons—which are stabilized by base-pairing and well-isolated in the downfield region—from 600 MHz to 1.3 GHz for a 14-mer hairpin fragment derived from an HIV-1 Vif-targeting RNA aptamer [9], which incorporates 2’-fluorine substitutions on the ribose to enhance nuclease resistance for pharmaceutical development. While the ^1^*J*_NH_ values of our sample were almost constant from 600 to 900 MHz, a significantly different value was observed exclusively at 1.3 GHz. This difference represents the direct detection of magnetic field-induced alignment without any alignment media. This work demonstrates that 1.3 GHz NMR successfully captures field-induced RDCs that were elusive at lower magnetic fields, highlighting ultra-high-field NMR as an indispensable tool for the media-free structural analysis of short nucleic acids.

## Results and Discussion

To gain detailed structural insights into the 14-mer RNA used in this study, we first performed comprehensive NMR signal assignments for three-dimensional structure determination. Experimental details are provided in the Supporting Information. Analysis of COSY, TOCSY, NOESY, ^1^H,^13^C HSQC, ^1^H,^19^F HSQC, and ^1^H,^19^F HOESY spectra yielded complete assignments for all non-exchangeable protons, imino protons, and the 2’-F nuclei. The three-dimensional structure of the RNA was determined using NOE-derived distance restraints, hydrogen-bonding restraints, and dihedral angle restraints (Fig. 1). As shown in Fig. 1, the 14-mer RNA forms an A-form stem with a U6–A7–G8–A9 hairpin loop. The NOESY spectra indicated that the G8 base, located at the center of the loop, is partially stacked with the adjacent A7 and A9 bases to form an ordered local conformation. Meanwhile, a limited number of NOE cross-peaks within the loop region and the line broadening of the U6 ^19^F signal suggest the structural flexibility of the loop. This U6–A7–G8–A9 sequence is considered a crucial recognition interface when this RNA binds to protein complexes containing HIV-1 Vif. [9].

**Figure 1.**
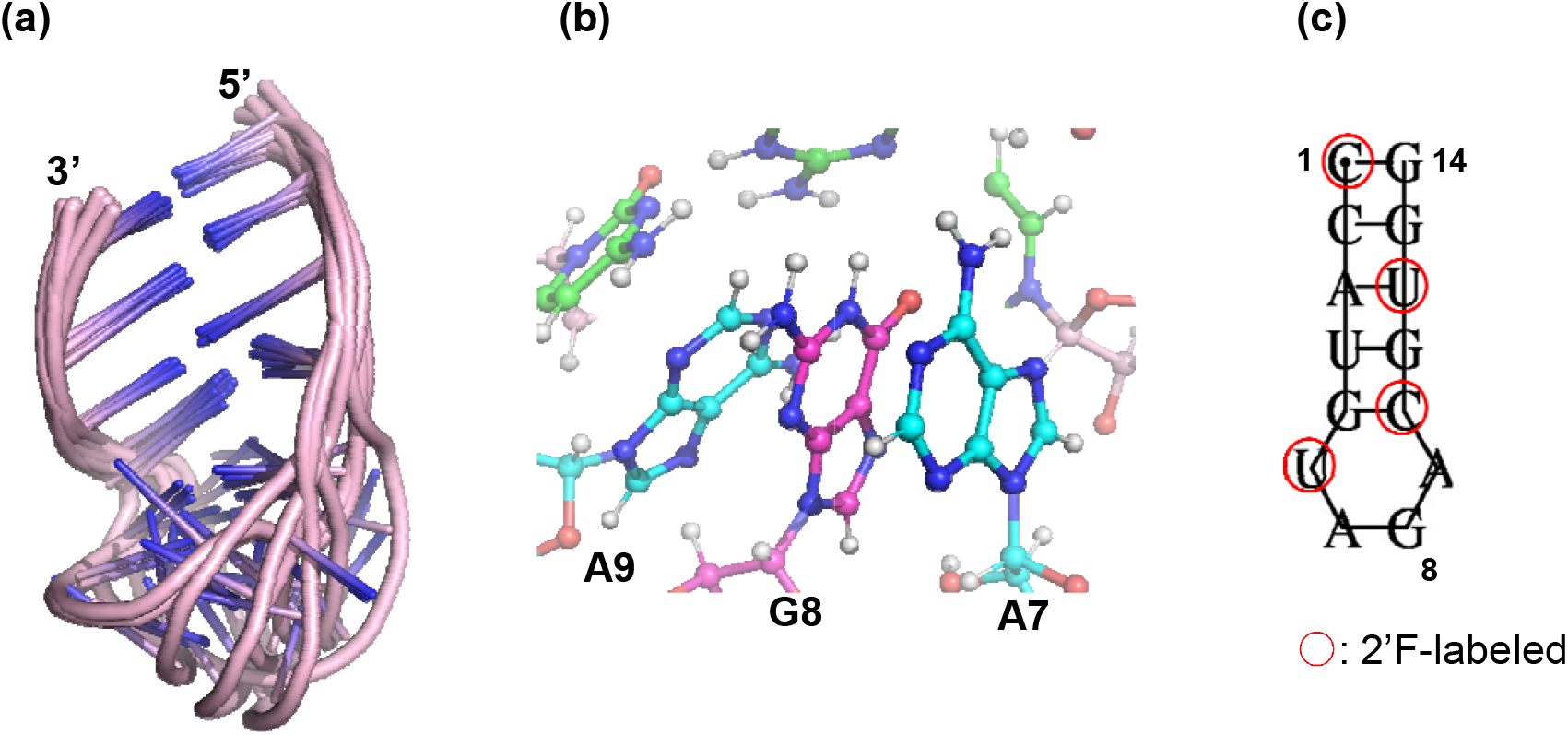
NMR structure of the RNA hairpin and local structural features of G8. (a) Ensemble of the 10 lowest-energy structures calculated with NOE and hydrogen-bonding restraints, showing an A-form helical stem capped by a UAGA tetraloop. (b) Close-up view of the A7–G8–A9 segment. The aromatic ring of G8 is sandwiched between adjacent bases, consistent with partial stacking interactions supported by NOESY data. (c) Secondary structure of the RNA hairpin, indicating base-pairing interactions in the stem region.

To independently confirm the determined RNA structure, we utilized residual dipolar couplings (RDCs) obtained from magnetic field-induced alignment. Therefore, NMR measurements were performed at multiple field strengths of 600, 700, 800, 900, and 1300 MHz. Specifically, scalar coupling constants (^1^*J*_NH_) were measured from ^1^H,^15^N SOFAST-HMQC spectra [10] without ^15^N decoupling during data acquisition (see Supporting Information for technical details). 1D slices of ^1^H–^15^N correlation peaks at each magnetic field strength were compared for two representative residues exhibiting different field-dependent behaviors (Fig. 2). Notably, the horizontal axis is displayed in Hz to enable a direct comparison across the different magnetic fields, where the splitting corresponds to the scalar coupling constant (^1^*J*_NH_). For G14 (Fig. 2A), located in the stable stem region (Fig. 1), the TROSY component became sharper with increasing magnetic field strength, leading to enhanced signal intensity.[11] At 1.3 GHz, the TROSY effect yielded peaks with the highest resolution and sensitivity. In terms of transverse relaxation (*T*_2_), the TROSY effect is theoretically optimal around 900 MHz. Nevertheless, the intrinsic gains in signal strength at 1.3 GHz compensated for this relaxation disadvantage, yielding a higher apparent sensitivity than at 900 MHz.[8] Quantitative evaluation of the magnetic field dependence of ^1^*J*_NH_ revealed no significant changes between 600 and 900 MHz, whereas a clear deviation was detected only at 1.3 GHz (Fig. 2A). Furthermore, a plot of ^1^*J*_NH_ values demonstrated that similar changes emerged exclusively at 1.3 GHz for all observed residues excluding G8 (Fig. 3). These residues are located within the stem region of the RNA (Fig. 1). Given that the ^1^*J*_NH_ values remained virtually constant between 600 and 900 MHz, the 600 MHz data were taken as the isotropic scalar coupling constant (Figs 2A and 3). The experimental RDC values at 1.3 GHz were then defined as the difference in ^1^*J*_NH_ between 600 MHz and 1.3 GHz. To validate these values, we evaluated their agreement with the NOE-derived structure by calculating the Q-factor using the PALES program [12]. The low Q-factor (0.28) obtained for these residues confirms that the observed variations at 1.3 GHz are indeed structural RDCs.

**Figure 2.**
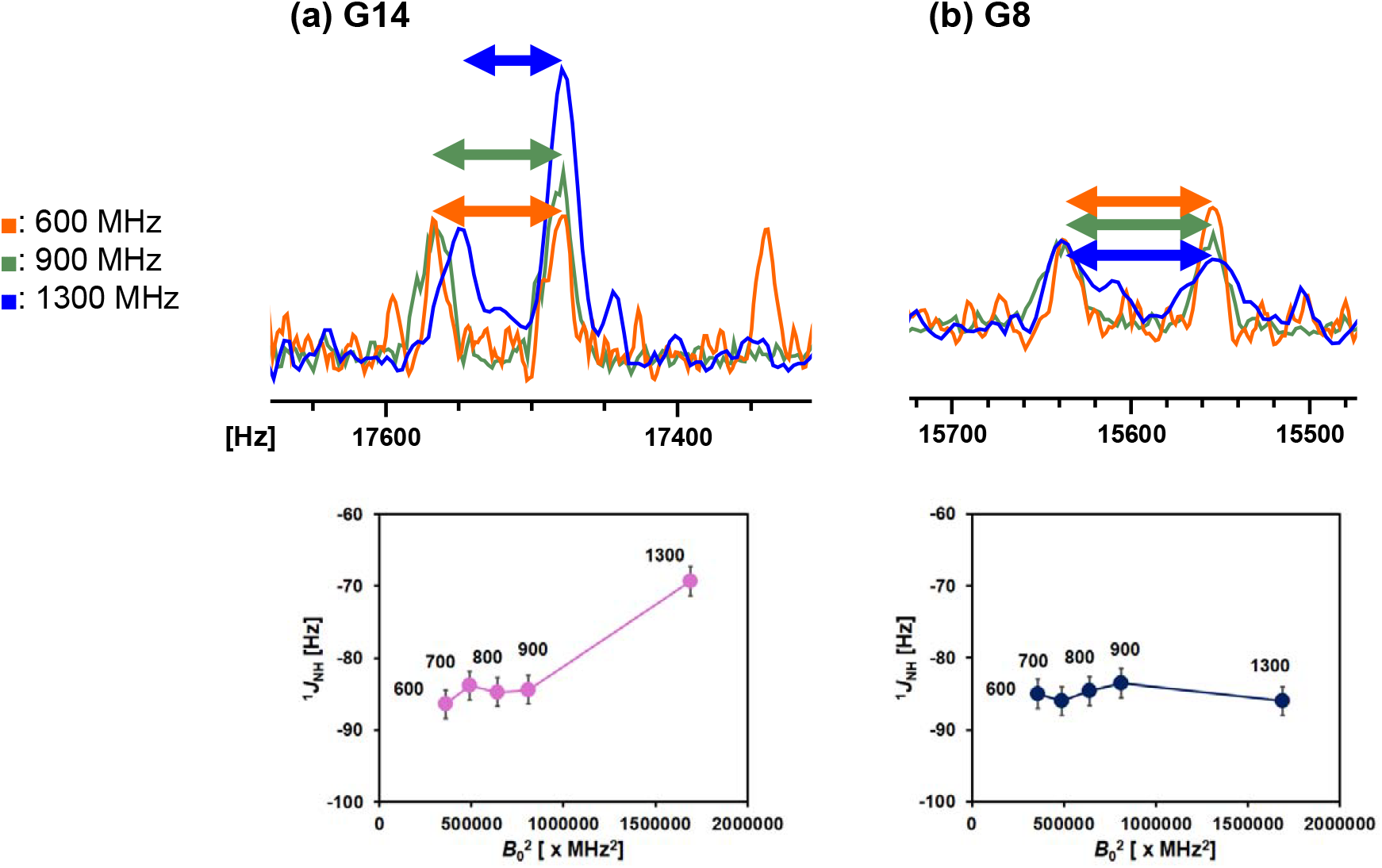
Field-dependent spectral changes in representative residues. (a) (upper) One-dimensional slices of ^1^H,^15^N SOFAST-HMQC spectra for a representative base-paired residue in the stem region (e.g., G14) at different magnetic field strengths (orange: 600 MHz; green: 900 MHz; blue: 1.3 GHz). The horizontal axis is shown in Hz. The TROSY peak top positions at 600 and 900 MHz were individually aligned to the peak top of the TROSY component at 1.3 GHz. The ^1^*J*_NH_ values at each field strength are indicated by color-coded double-headed arrows corresponding to each spectrum. A pronounced change in ^1^*J*_NH_ is observed at 1.3 GHz. For comparison, the intensities of the anti-TROSY components were normalized across different field strengths. (lower) Magnetic field dependence of ^1^*J*_NH_, showing a marked increase at 1.3 GHz. (b) (upper) Corresponding data for G8 located in the loop region. (lower) Magnetic field dependence of ^1^*J*_NH_ for G8, showing no significant variation across the measured field range.

**Figure 3.**
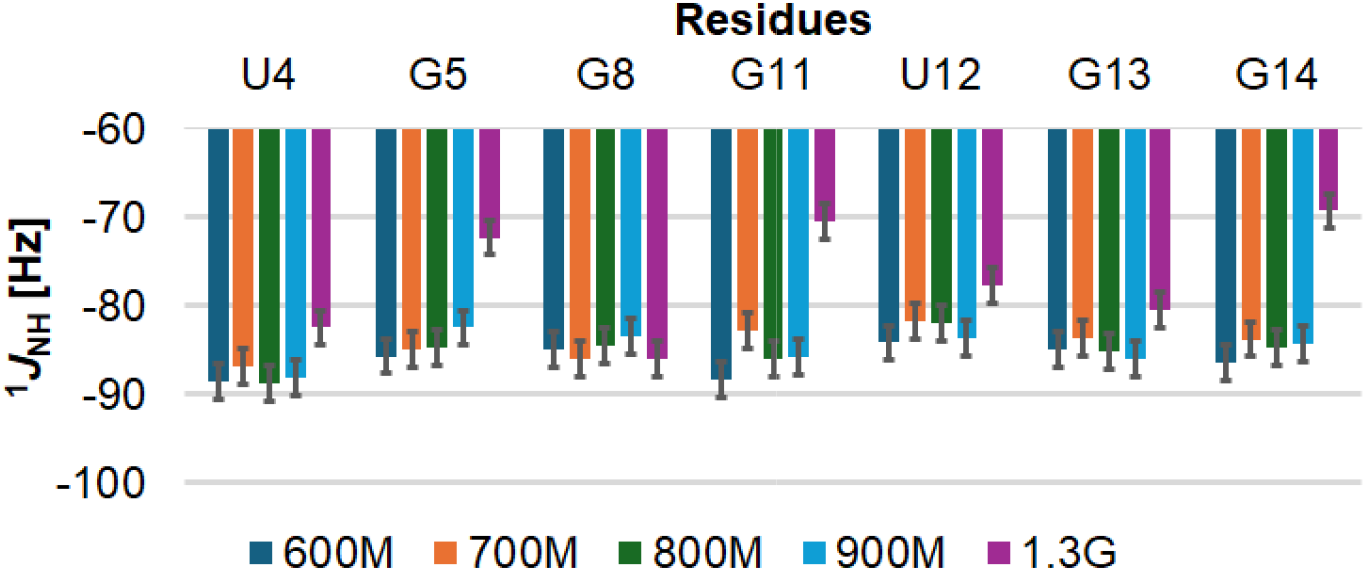
Field dependence of residual dipolar couplings. Residue-resolved changes in ^1^H–^15^N scalar couplings (^1^*J*_NH_) at different magnetic field strengths. At 600–900 MHz, ^1^*J*_NH_ values remain within experimental uncertainty for all residues. In contrast, multiple residues exhibit substantial changes in ^1^*J*_NH_ at 1.3 GHz, indicating that RDC contributions become detectable at this field strength. Notably, G8 shows negligible variation in ^1^*J*_NH_ across all fields.

Theoretically, the degree of field-induced alignment is governed by the magnetic susceptibility anisotropy (Δ*χ*), where the resulting alignment energy is proportional to the square of the field strength (*B*_0_^2^). This alignment inherently competes with the thermal energy (*kT*). Unfortunately, for an RNA of this molecular size, both the degree of alignment and the detection sensitivity are insufficient at 600–900 MHz to observe the RDCs. The 1.3 GHz environment successfully overcomes this limitation, driving a synergistic effect that scales as 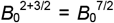 through the combination of alignment 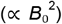 and sensitivity 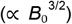. This physical scaling provides an approximate 15-fold and 4-fold increase in the overall RDC detectability at 1.3 GHz compared to 600 MHz [(1300/600)^7/2^ ≈ 15] and 900 MHz [(1300/900)^7/2^ ≈ 4], respectively. To verify whether the underlying 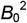 -dependence was obscured by experimental errors at lower fields, we simulated the expected 1.3 GHz values by extrapolating the lower-field datasets and their associated errors based on a 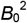 scaling (see Supporting Information for details). The actual experimental values obtained at 1.3 GHz fell within these predicted error margins. This demonstrates that the lack of apparent changes at 600–900 MHz was simply because the RDC values in this regime were smaller than the intrinsic uncertainty, rather than the absence of field-induced alignment. Therefore, our study represents the first instance of achieving observation of alignment for short RNA of this size using a 1.3 GHz spectrometer.

The imino proton of G8 was observable despite the absence of canonical hydrogen bonding within the loop. An unpaired imino proton within a flexible loop typically undergoes rapid chemical exchange with water, leading to signal loss. The visibility of the G8 resonance suggests that the stacking with adjacent A7 and A9 bases may restrict solvent accessibility (Fig. 1). Although the present UAGA tetraloop is structurally distinct from the structured internal motifs or rigid ligand-binding pockets previously documented [13, 14], these studies demonstrate that base packing can sequester labile imino protons from the bulk solvent even in the absence of canonical hydrogen bonds. Despite this protective environment allowing the observation of the G8 imino proton, no RDC was detected across all measured field strengths (Figs. 2 and 3). As noted above, this loop region undergoes local conformational dynamics that cause spatial averaging of the angular distribution, driving the observed RDC values toward zero. [1, 2, 15] Thus, G8 exemplifies a residue that is structurally ordered enough to retain its imino proton signal, yet dynamically flexible enough to suppress any detectable alignment. Intriguingly, G8 exhibited an unexpected magnetic field dependence in the relative intensities of its TROSY and anti-TROSY components (Fig. 2B). Normally, the TROSY effect causes the lower-frequency component along the ^1^H axis to be sharper and more intense.[11] For G8, however, the lower-frequency component broadened with increasing magnetic field, and its intensity was inverted relative to the higher-frequency component at 1.3 GHz. The transverse relaxation rates (*R*_2_) of the TROSY 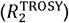 and anti-TROSY 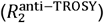 components along the ^1^H axis are expressed as follows:

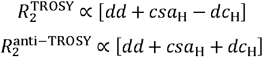

where *dd* and *csa*_H_ represent the independent dipole-dipole (DD) and ^1^H chemical shift anisotropy (CSA) contributions, respectively. The *dc*_H_ term denotes the DD/CSA cross-correlation, which is proportional to the geometric factor (3 cos^2^ *θ* −1)/2, where *θ* is the relative angle between the N-H dipolar vector and the principal axis of the ^1^H CSA tensor. For typical imino groups involved in standard base-pairing, theoretical studies have established that the N-H vector and the ^1^H CSA principal axis are nearly parallel (*θ* ≈ 0°), thereby maintaining a positive *dc*_H_ contribution. [16, 17] Consequently, the TROSY component is sharper and more intense than the anti-TROSY component. For G8, however, the observed line-width inversion directly reflects a change in the sign of the *dc*_H_ term. This negative *dc*_H_ contribution indicates that the angle *θ* in G8 is substantially altered, to the extent that it flips the sign of the geometric factor. This atypical angular relationship strongly suggests an unusual local electronic distribution around the G8 imino proton. These findings provide a clear physical rationale for the NOESY-derived model, in which the G8 base maintains partial stacking with the adjacent A7 and A9 bases (Fig. 1). This restricted local geometry is expected to distort the electronic distribution around the G8 imino proton relative to that in a standard base-pair. This structural dualism of G8—balancing dynamic loop flexibility with ordered local packing—likely optimizes the tetraloop conformation for its specific functional role as a recognition interface. [9]

More broadly, our findings highlight the indispensable role of 1.3 GHz NMR in structural biology. The ultra-high magnetic field environment not only overcomes the physical limitations of detecting subtle self-alignment in short, fast-tumbling molecules but also provides the critical resolution required to capture local geometric and electronic anomalies. This approach opens up new avenues for elucidating the precise structures and physicochemical properties of functional sites within a wide range of target RNAs.

## Acknowledgements

We are deeply grateful to Drs. Kümmerle and Mayzel (Bruker BioSpin), Dr. Sato (Bruker Japan), and Drs. Cheong, Lee, and Ryu (KBSI, Korea) for providing access to the UHF NMR spectrometers and for their generous support. We would also like to thank S. Yasuda for her secretarial assistance. We would also like to thank S. Yasuda, A. Yokooku, and A. Sekiguchi for their secretarial assistance. The research was supported in part by “NMR Platform” supported by the Ministry of Education, Culture, Sports, Science and Technology (MEXT), Grant Number JPMXS0450100021, from Ministry of Education, Culture, Sports, Science and Technology, Japan.

## Data Availability Statement

The data that support the findings of this study are available in the supplementary material of this article.

