## Supporting Information for "Expanding the Frontiers of Structural Analysis in Short RNAs by Ultra-High Field 1.3 GHz NMR"

### 1. Experimental Details

#### 1.1. Sample

The RNA sample was purchased from Hokkaido System Science Co., Ltd. The sample was dissolved in nuclease-free ultrapure water and annealed by heating at 363 K for 5 min and snap-cooling on ice. The sample was then buffer-exchanged into the NMR buffer (20 mM sodium phosphate buffer pH 6.5 containing 50 mM NaCl) using Vivaspin2 centrifugal concentrators (MWCO 2kDa; Sartorius, Göttingen, Germany). Quantification via UV absorbance at 260 nm determined the sample concentration to be 0.3 mM. The sample was analyzed by native PAGE to confirm the hairpin structure.

#### 1.2. NMR experiments

NMR experiments were performed on Bruker Avance III HD spectrometers operating at 600, 700, and 800 MHz equipped with 5 mm cryogenic probes (QCI for 600 MHz and TXO for 700 and 800 MHz), a Bruker Avance NEO 900 MHz spectrometer equipped with a 5 mm cryogenic TCI-F probe, and a Bruker Avance NEO 1.3 GHz spectrometer equipped with a 5 mm cryogenic TXO probe. The 600–900 MHz spectrometers were located at RIKEN (https://www.ynmr.riken.jp), while the 1.3 GHz measurements were performed on a pre-commercial prototype system provided by Bruker (Fällanden, Switzerland). All experiments were conducted at 283 K unless otherwise noted.

SOFAST-HMQC experiments were performed as described by Schanda and Brutscher [1]. Selective $^{1}$H excitation was centered at 12.25 ppm with a bandwidth of 3.5 ppm. $^{15}$N decoupling during acquisition was achieved using a GARP composite pulse scheme. For $^{1}J_{NH}$ measurements, the GARP decoupling power was set to 0 W. In the $^{1}$H direct dimension, 6k–13k complex data points were acquired without $^{15}$N decoupling, covering a spectral width of 20 ppm centered at 4.7 ppm. In the $^{15}$N indirect dimension, 32 complex points were recorded over 30 ppm centered at 155 ppm. When $^{15}$N decoupling was applied, 1024 complex points were collected in the $^{1}$H dimension, while the $^{15}$N dimension was unchanged. The relaxation delay was set to 100 ms, and 5k–8k scans were accumulated depending on the signal sensitivities. The spectral resolution in the $^1$H dimension was sufficient to resolve the splitting between TROSY and anti-TROSY components for accurate estimation of $^1J_{NH}$ values. The experimental resolution in the $^1$H dimension was approximately 4 Hz. This value was taken as an estimate of the uncertainty in the determination of $^1J_{NH}$. To statistically verify the presence of magnetic field-induced alignment within the lower-field datasets (600, 700, 800, and 900 MHz), an extrapolation analysis was performed. To account for measurement uncertainty, a Monte Carlo simulation with 128 iterations was executed. In each iteration, synthetic $^1J_{NH}$ values for the lower fields were generated by adding random noise drawn from a normal distribution with a standard deviation (SD) of 4 Hz to the experimentally measured values. For each simulated dataset, a linear regression was performed against $B_0^2$ to extrapolate the $^1J_{NH}$ value at 1.3 GHz. After 128 trials, the mean and the standard deviation of the predicted 1.3 GHz values were calculated.

For chemical shift assignment, conventional 2D NMR experiments were recorded, including $^1$H,$^1$H COSY, TOCSY, and NOESY, as well as $^1$H,$^{13}$C HSQC, $^1$H,$^{19}$F HSQC, and $^1$H,$^{19}$F HOESY spectra. $^1$H,$^1$H NOESY spectra were acquired with mixing times of 100 and 400 ms. The $^1$H,$^{19}$F HOESY experiment was recorded with a mixing time of 400 ms. $^1$H,$^1$H NOESY spectra were measured at 278, 283, and 288 K.

### 1.3. Structure determination

#### Structure Calculation Using XPLOR-NIH

Structure calculations were performed using XPLOR-NIH [2]x. Initial structures were generated from randomized conformations, followed by structure calculations using distance restraints derived from NOESY spectra, hydrogen-bond restraints for base-paired residues, and dihedral angle restraints. Simulated annealing was performed starting at 3500 K for 50,000 steps, followed by a cooling schedule from 3500 K to 25 K over 240 temperature steps (12.5 K increments), with 500 steps at each temperature, resulting in a total of 120,000 cooling steps. A total of 50 structures were generated, and the lowest-energy structures were selected based on total energy and restraint violations. These structures were used to evaluate the consistency between the experimentally observed RDCs and the calculated structures using PALES [3].

#### AMBER Refinement

The selected 50 structures from XPLOR-NIH were further refined using the AMBER simulation package (AMBER22, Case, D.A.; et al., University of California, San Francisco). Topology and coordinate files were generated using the tleap module with the leaprc.RNA.OL3 force field. To incorporate 2′-fluoro modifications, the 2′-OH group was replaced by fluorine by removing the HO2′ atom and substituting the O2′ atom with fluorine, followed by appropriate modification of atom types and residue names. Partial atomic charges for the modified nucleotides were assigned using the AM1-BCC method as implemented in antechamber, assuming a net charge of −1 per nucleotide [4]. Bond orders in intermediate MOL2 files were manually corrected prior to charge calculation. Experimental restraints, including NOE-derived distance restraints, hydrogen-bond restraints, and dihedral angle restraints, were converted into AMBER restraint (RST) format and applied during refinement. Energy minimization was performed, followed by simulated annealing using SANDER/PMEMD. Annealing was carried out from 300 K to 0 K under restraint conditions to relieve local structural distortions while preserving experimentally derived constraints. The final structural ensemble consists of the 10 lowest-energy structures selected based on total energy and restraint violations after AMBER refinement.

2. Supplementary Figures

Figure S1. Comparison of $^{1}H$,$^{15}N$ SOFAST-HMQC spectra recorded at 1.3 GHz with and without $^{15}N$ decoupling. (a) Spectrum recorded with $^{15}N$ decoupling. (b) Spectrum recorded without $^{15}N$ decoupling. Resonance assignments are indicated in both spectra. In the absence of $^{15}N$ decoupling (b), clear doublet splittings are observed for each cross peak, allowing direct measurement of $^{1}J_{NH}$ values.

Figure S2. $^{19}F$ 1D NMR spectrum of the RNA containing site-specific 2′-fluoro substitutions, recorded with $^{1}H$ decoupling.

All 3 fluorinated residues except U6 exhibit sharp signals with similar chemical shifts. In contrast, the signal corresponding to U6 is significantly broadened and shifted downfield relative to the other residues. This distinct behavior indicates increased local dynamics and a different chemical environment for U6 in the loop region.

Figure S3. Residue-resolved comparison of $^{1}H$,$^{15}N$ SOFAST-HMQC 1D slices at different magnetic field strengths. Each panel corresponds to an individual residue, allowing comparison across all residues.

One-dimensional slices for individual residues are shown at 600 (orange), 700 (yellow), 800 (purple), 900 MHz (green), and 1.3 GHz (blue). The horizontal axis is given in Hz. For each residue, spectra at different fields were aligned to the peak top of the TROSY component at 1.3 GHz. The dashed blue vertical line indicates the position of the TROSY peak at 1.3 GHz, while the solid blue vertical line indicates the corresponding anti-TROSY peak at 1.3 GHz. The solid orange vertical line represents the anti-TROSY peak position at 600 MHz. The difference in $^{1}J_{NH}$ between 1.3 GHz and 600 MHz is indicated for each residue and was used as the RDC value.

3. Supplementary Tables

Table S1. Residue-resolved $^{1}J_{NH}$ values measured at different magnetic field strengths and the extrapolated 1.3 GHz values based on the 600-900 MHz datasets. All values are given in Hz.

| Residue | 600M | 700M | 800M | 900M | 1.3G | 1.3G (extrapolated) |
|---|---|---|---|---|---|---|
| **U4** | −88.6 | −86.9 | −88.7 | −88.2 | −82.5 | −89.58 ± 12.98 |
| **G5** | −85.7 | −84.9 | −84.7 | −82.5 | −72.3 | −76.62 ± 12.79 |
| **G8** | −85.0 | −86.0 | −84.6 | −83.5 | −86.0 | −81.25 ± 13.59 |
| **G11** | −88.3 | −82.8 | −86.0 | −85.8 | −70.4 | −84.97 ± 13.04 |
| **U12** | −84.2 | −81.8 | −81.9 | −83.7 | −77.7 | −82.59 ± 13.61 |
| **G13** | −85.0 | −83.7 | −85.1 | −86.0 | −80.5 | −88.90 ± 13.62 |
| **G14** | −86.4 | −83.8 | −84.7 | −84.4 | −69.3 | −80.87 ± 12.15 |

Table S2. Structure statistics for the RNA ensemble

| | | | |
|---|---|---|---|
| Structure calculation[a] | | | |
| | Number of calculated structures: | | 50 |
| | Number of structures in final ensemble: | | 10 |
| Restraints | | | |
| | NOE restraints | | |
| | | Intra-residue: | 28 |
| | | Sequential: | 51 |
| | | Medium-range: | 3 |
| | | Long-range: | 8 |
| | Hydrogen bond restraints: | | 28 |
| | Dihedral angle restraints: | | 33 |
| Restraint violations | | | |
| | NOE violations > 0.3 Å: | | 0 |
| | NOE violations > 0.5 Å: | | 0 |
| | Dihedral violations > 5°: | | 0 |
| Structure quality[b] | | | |
| | RMSD to mean structure [Å] | | |
| | | Heavy atoms (all): | 3.09 ± 1.05 |
| | | Backbone atoms (all): | 3.12 ± 1.30 |
| | | Backbone atoms (stem): | 1.34 ± 0.53 |
| | | Backbone atoms (loop): | 5.38 ± 2.35 |

| | | |
|---|---|---|
| RDC statistics[c] | | |
| | Q factor: | 0.023 ± 0.016 |
| | RMS deviation (Hz): | 0.228 ± 0.165 |

[a]The number of calculated structures refers to the structures generated by XPLOR-NIH, whereas the final ensemble corresponds to the structures after AMBER refinement. [b]Restraint violations and structural quality metrics (RMSD) are reported for the AMBER-refined structures. [c]RDC statistics are reported as follows: both the Q factor and the RMS deviation (Hz) were calculated using PALES.

Figure S1

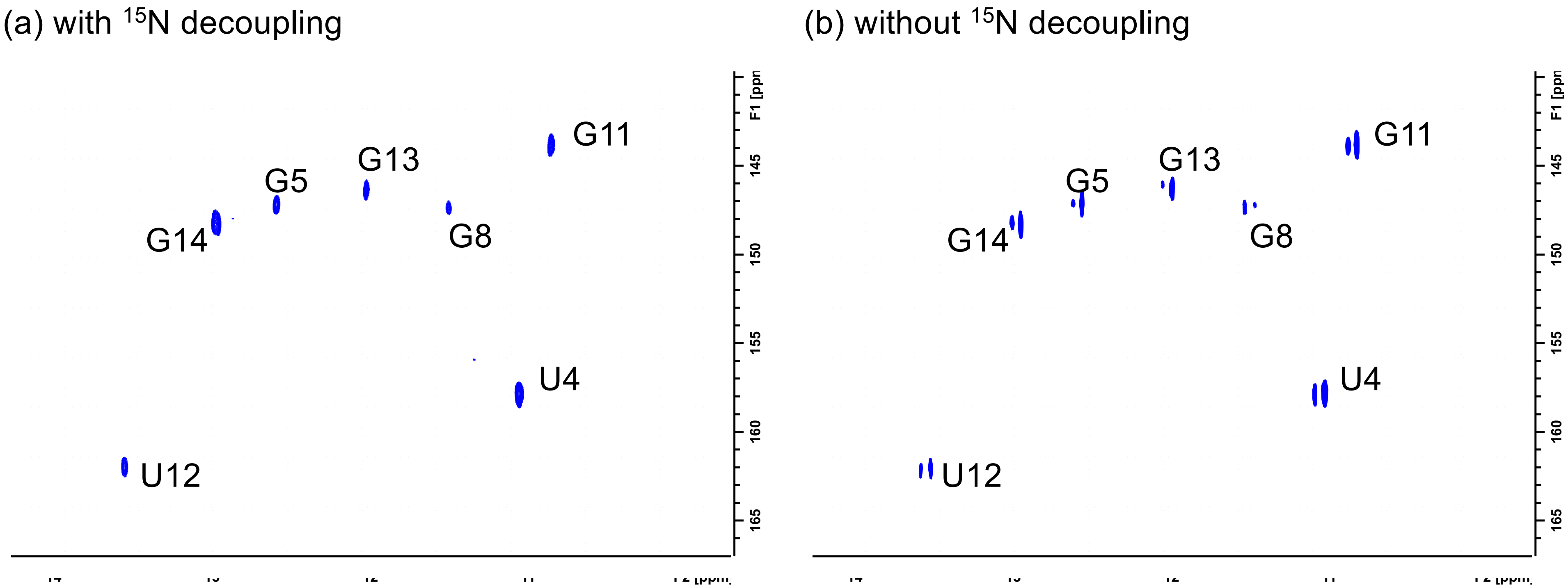
(a) with $^{15}$N decoupling
(b) without $^{15}$N decoupling
G11
G13
G5
G14
G8
U4
U12
F1 [ppm]
145
150
155
160
165
G11
G13
G5
G14
G8
U4
U12
F1 [ppm]
145
150
155
160
165

Figure S2

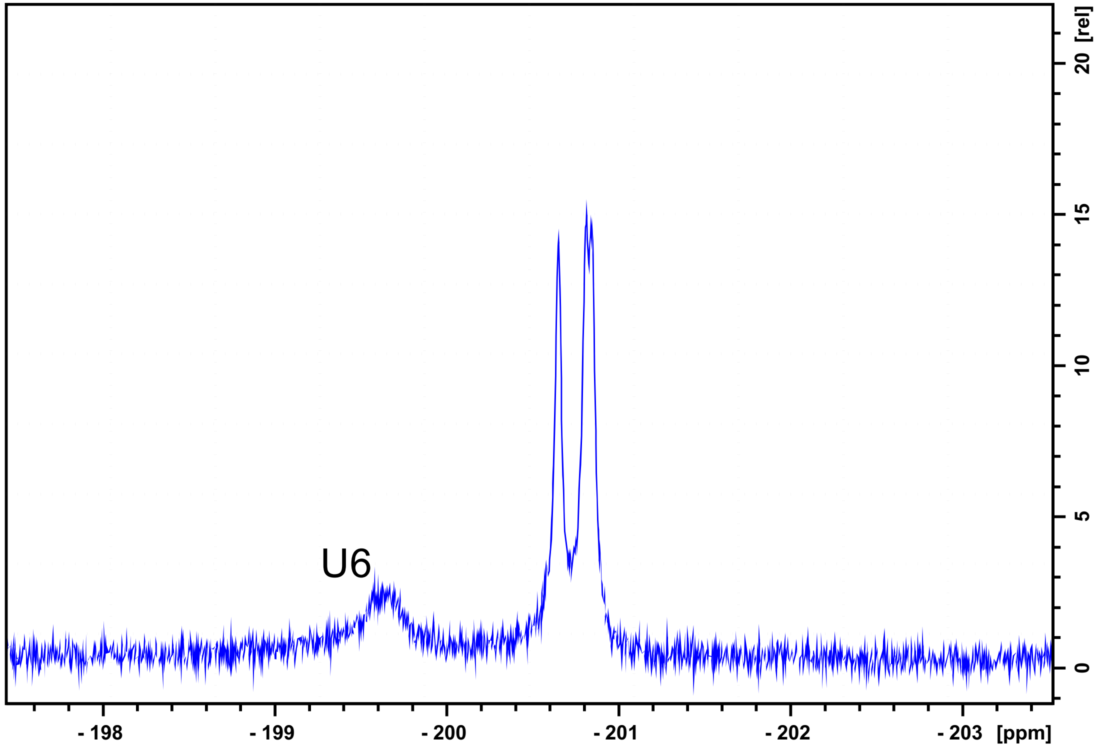

Figure S3

U4

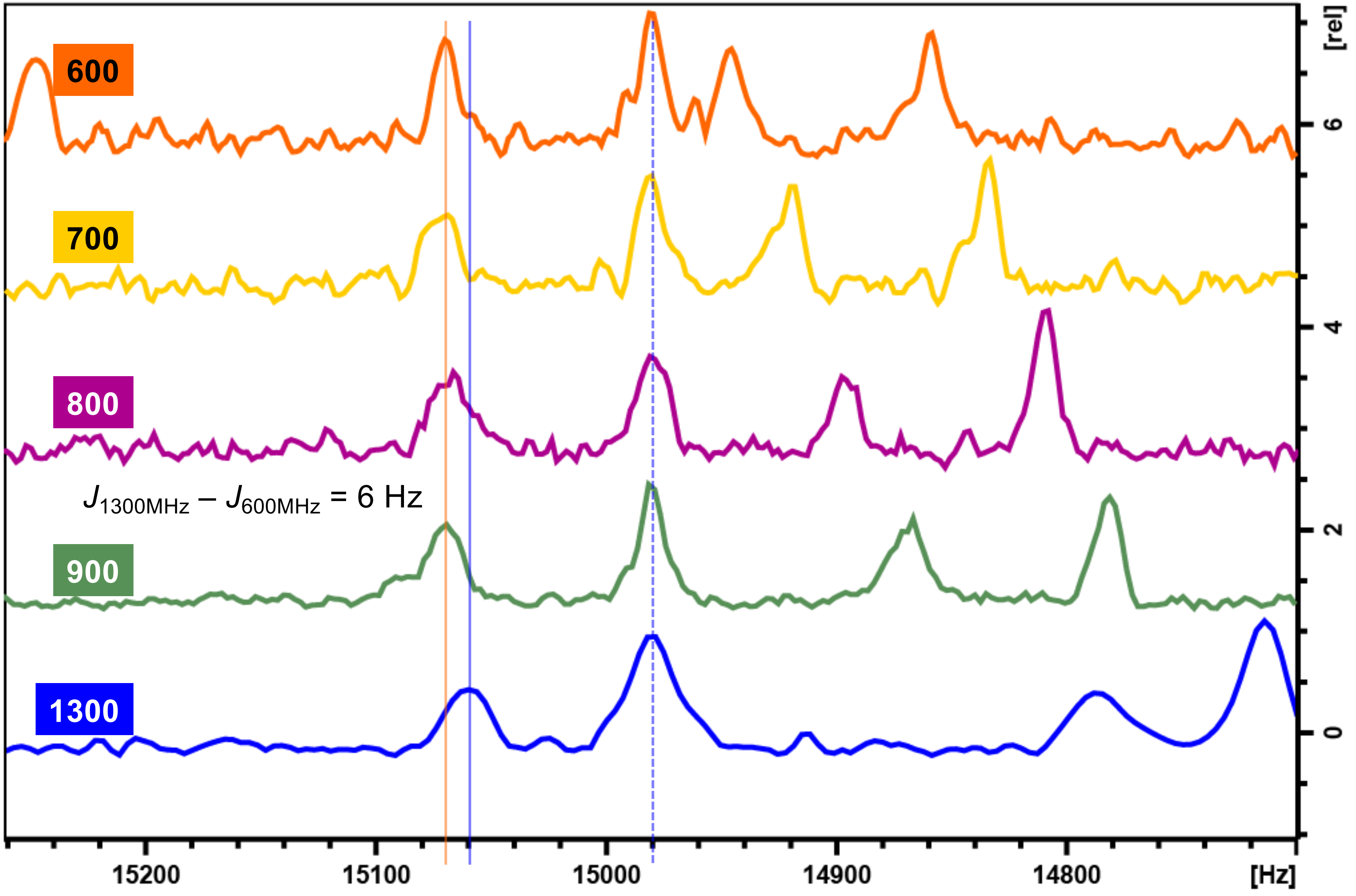

600
700
800
900
1300
$J_{1300MHz} - J_{600MHz}$ = 6 Hz
15200
15100
15000
14900
14800
[Hz]
[rel]
6
4
2
0

G5

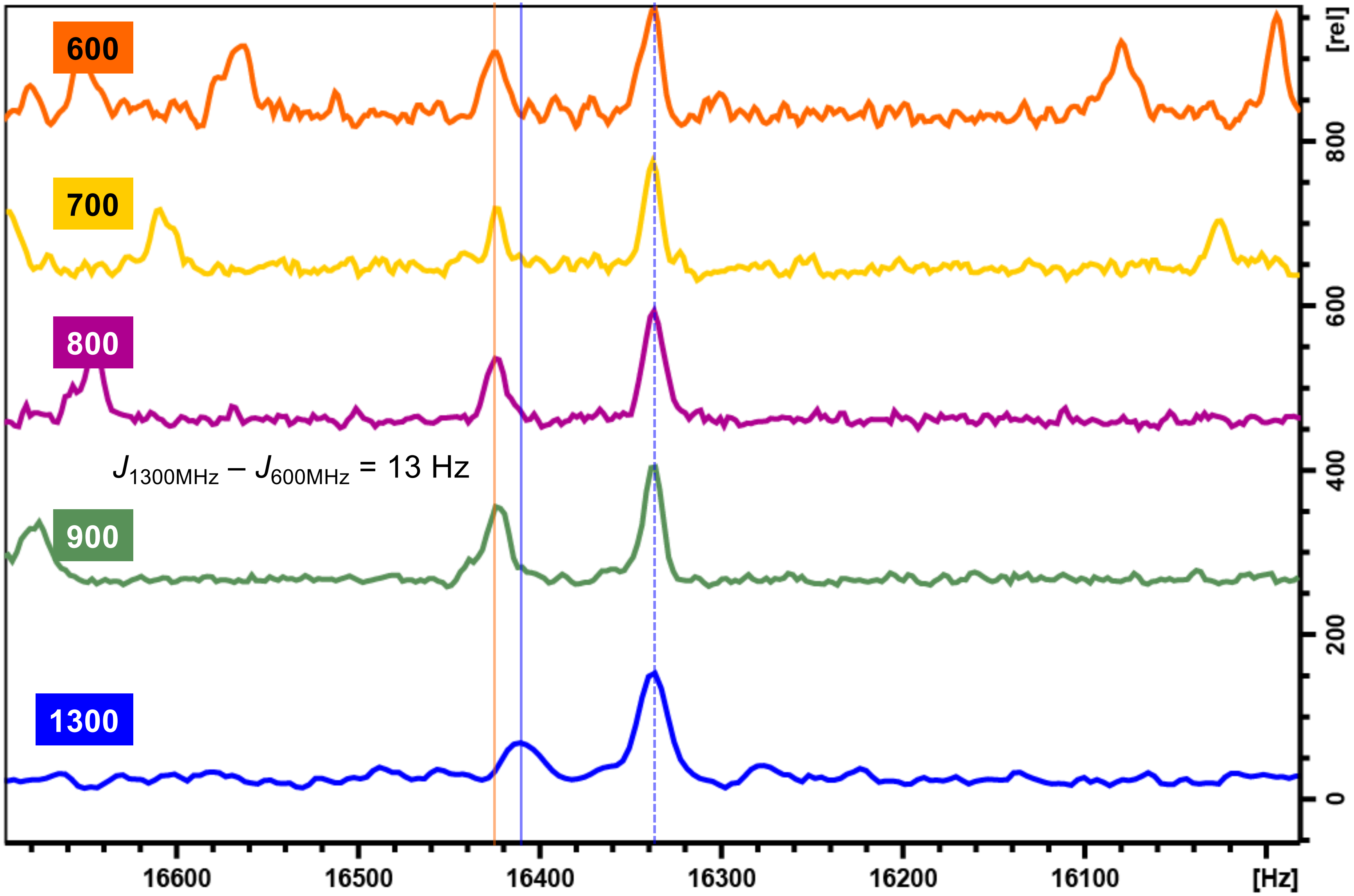
600
700
800
900
1300
$J_{1300MHz} - J_{600MHz}$ = 13 Hz
16600
16500
16400
16300
16200
16100
[Hz]
0
200
400
600
800
[rel]

G8

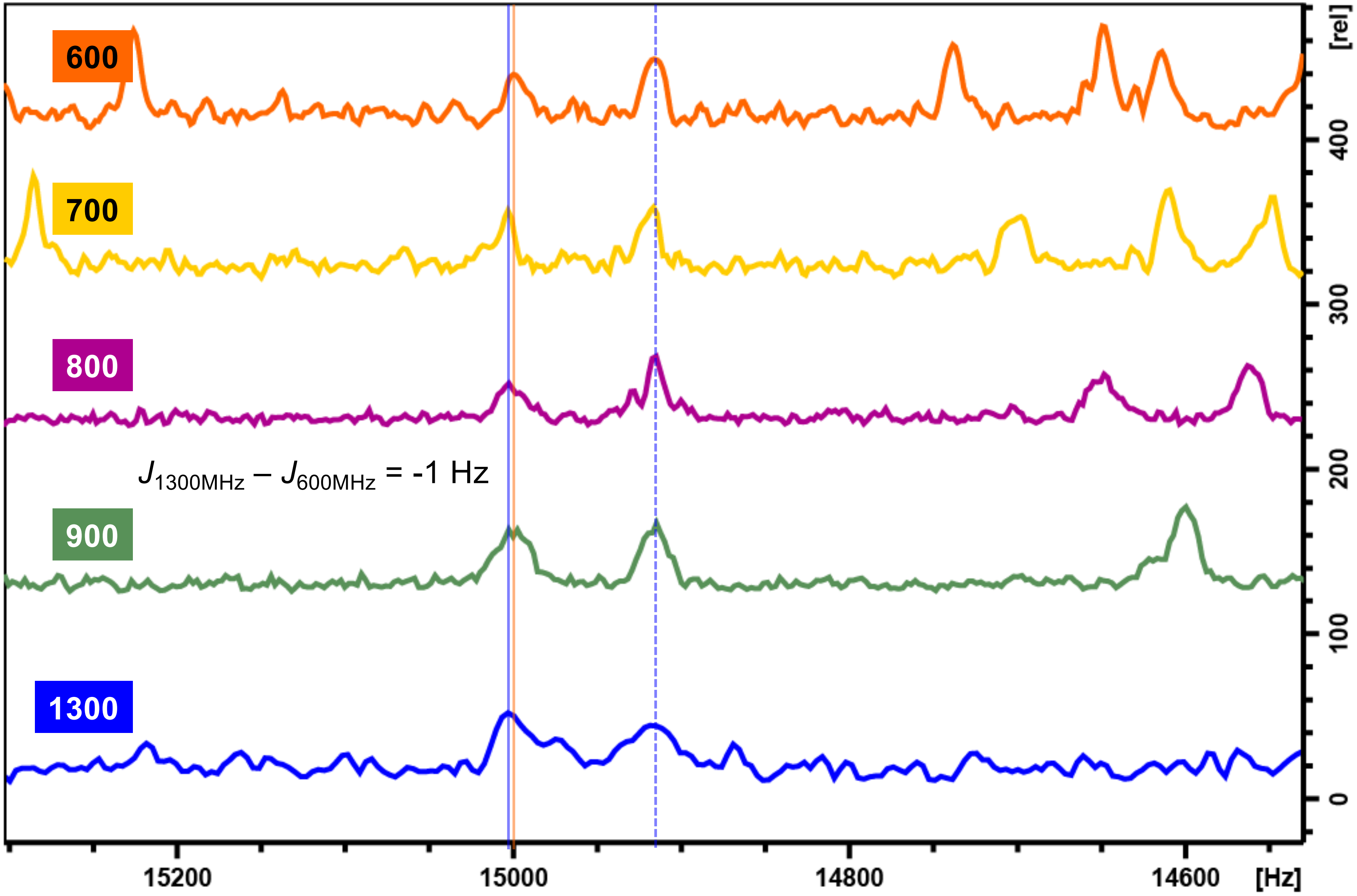
600
700
800
$J_{1300MHz} - J_{600MHz}$ = -1 Hz
900
1300
15200
15000
14800
14600
[Hz]
[rel]
0
100
200
300
400

G11

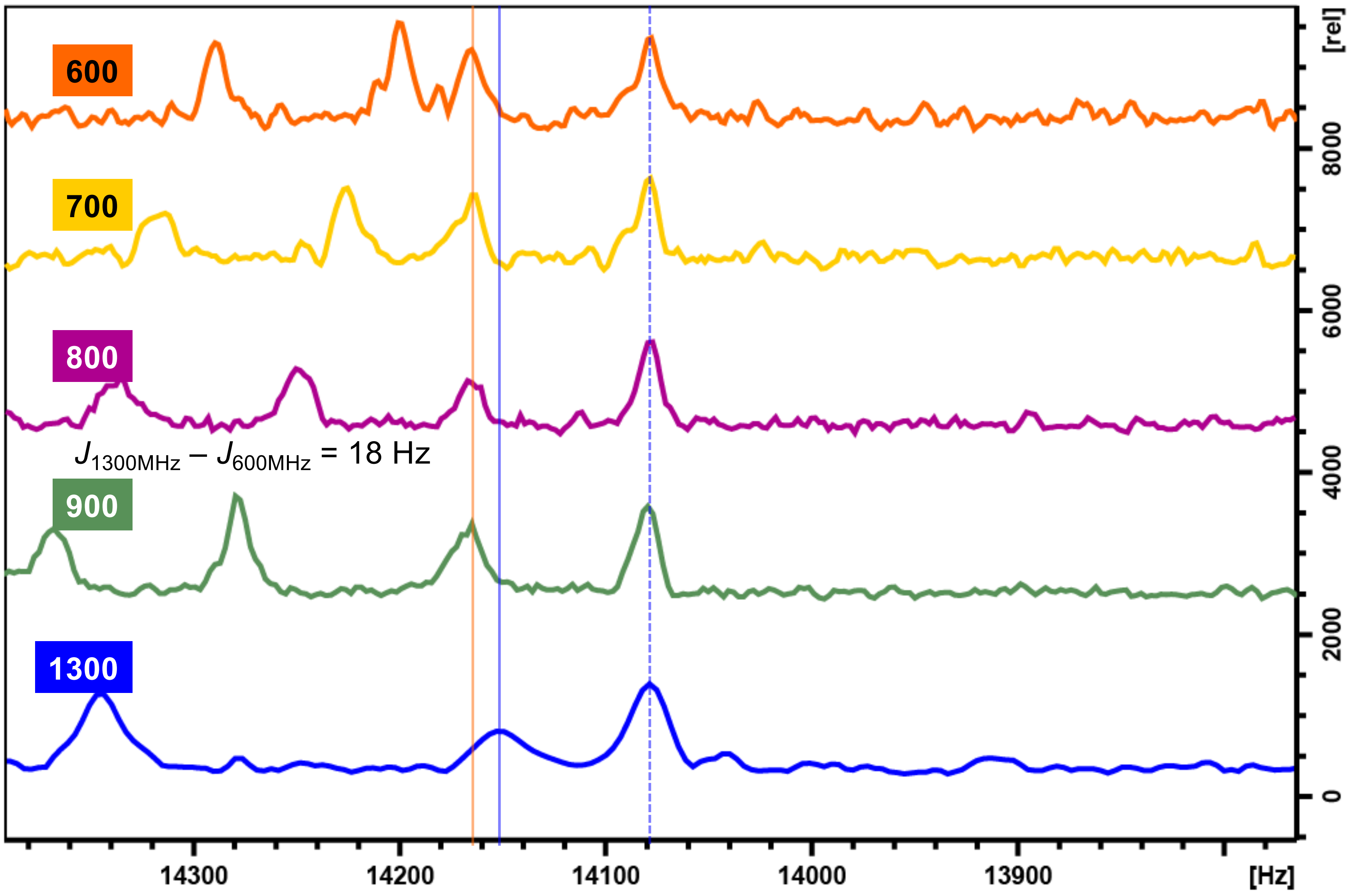

600
700
800
$J_{1300MHz} - J_{600MHz}$ = 18 Hz
900
1300
[rel]
8000
6000
4000
2000
0
14300
14200
14100
14000
13900
[Hz]

U12

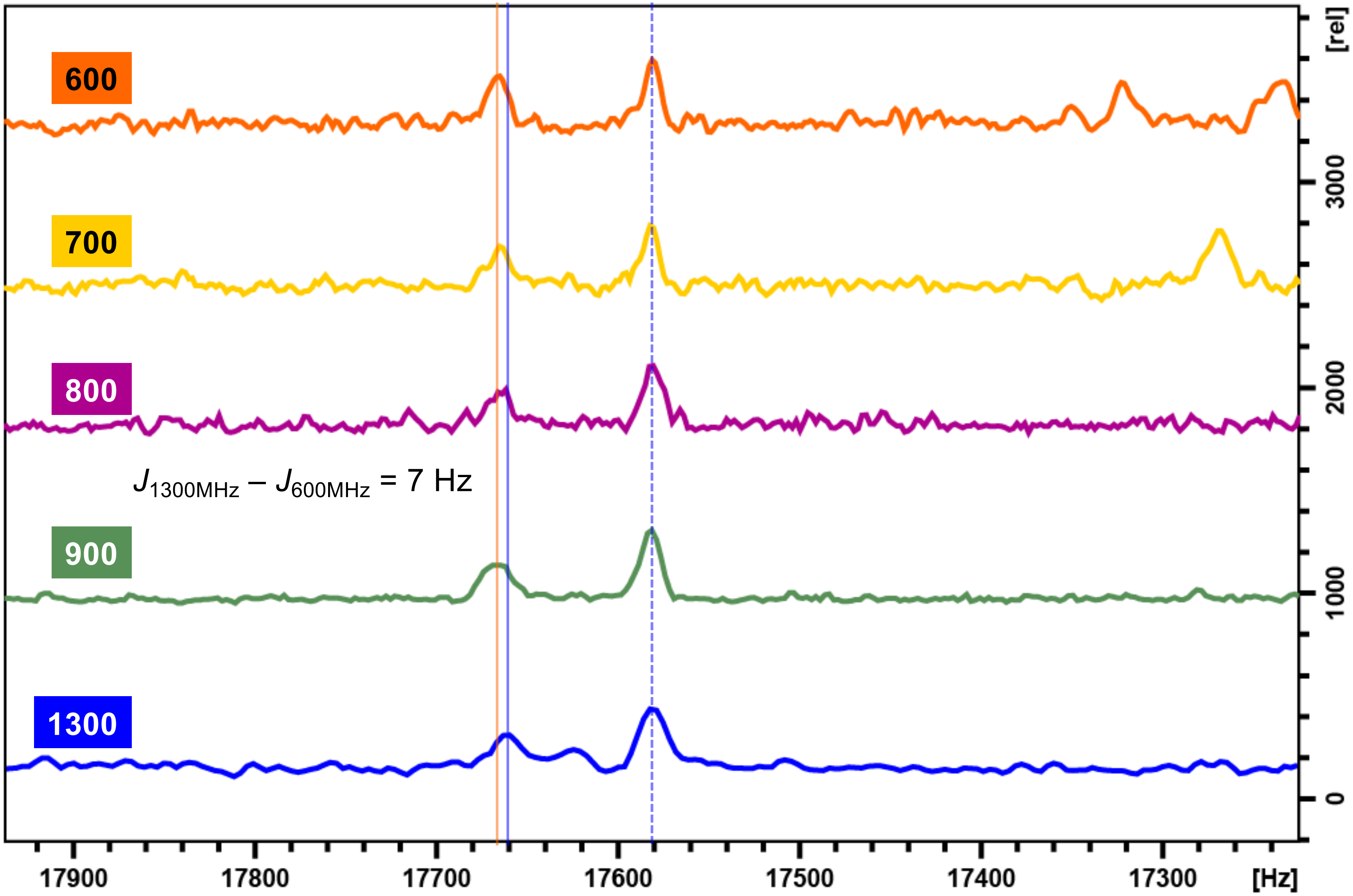

600
700
800
900
1300
$J_{1300MHz} - J_{600MHz}$ = 7 Hz
[rel]
3000
2000
1000
0
17900
17800
17700
17600
17500
17400
17300
[Hz]

G13

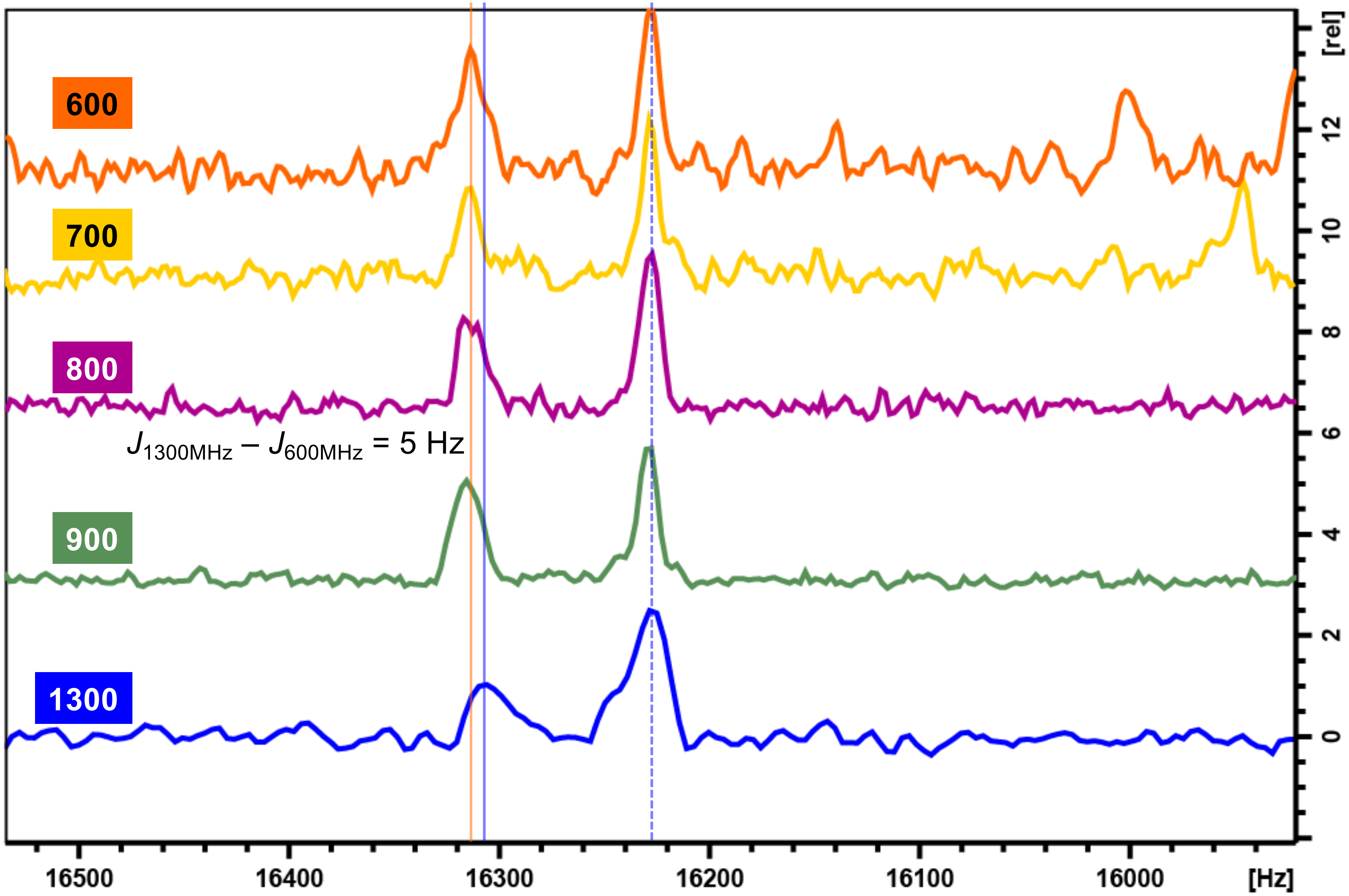
600
700
800
900
1300
$J_{1300MHz} - J_{600MHz}$ = 5 Hz
16500
16400
16300
16200
16100
16000
[Hz]
[rel]
12
10
8
6
4
2
0

G14

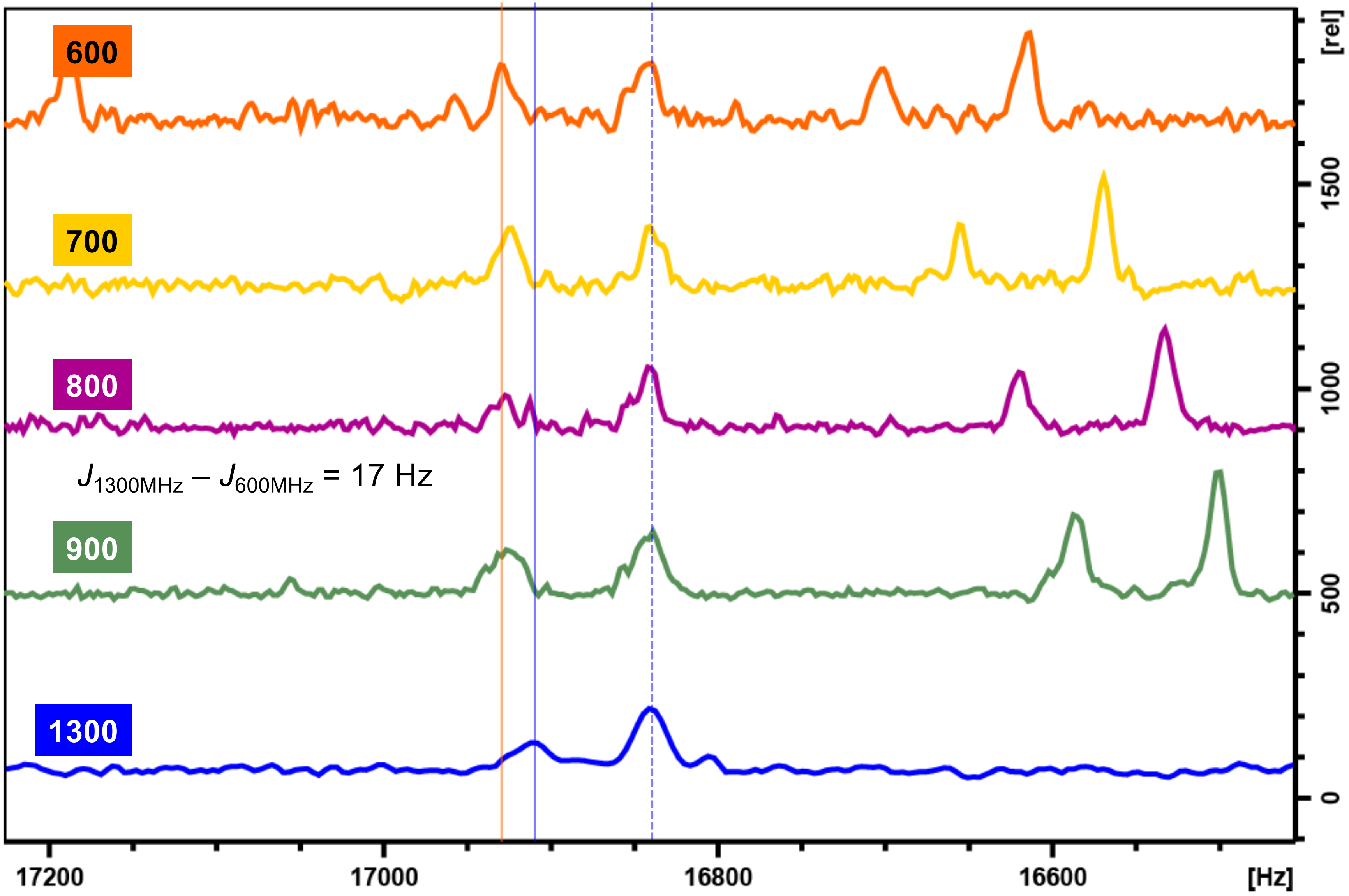

600
700
800
900
1300
$J_{1300MHz} - J_{600MHz}$ = 17 Hz
17200
17000
16800
16600
[Hz]
[rel]
1500
1000
500
0
